# Mad1’s ability to interact with Mad2 is essential to regulate and monitor meiotic synapsis in *C. elegans*

**DOI:** 10.1101/2021.05.14.444140

**Authors:** Alice Devigne, Needhi Bhalla

**Affiliations:** Department of Molecular, Cell and Developmental Biology, University of California, Santa Cruz, Santa Cruz, CA 95064

## Abstract

Meiotic homolog synapsis is essential to ensure accurate segregation of chromosomes during meiosis. In *C. elegans*, synapsis and a checkpoint that monitors synapsis relies on the spindle checkpoint components, Mad1 and Mad2, and Pairing Centers (PCs), cis-acting loci that interact with the nuclear envelope to mobilize chromosomes within the nucleus. Here, we show that mutations in some spindle checkpoint mutants affect PC movement early in meiotic prophase, consistent with a link between PC mobility and the regulation of synapsis. Further, we test what specific functions of Mad1 and Mad2 are required to regulate and monitor synapsis. We find that a mutation that abrogates Mad1’s localization to the nuclear periphery abolishes the synapsis checkpoint but has no effect on Mad2’s localization to the nuclear periphery or synapsis. By contrast, a mutation that prevents Mad1’s interaction with Mad2 abolishes the synapsis checkpoint, delays synapsis and fails to localize Mad2 to the nuclear periphery. These data indicate that Mad1’s primary role in regulating synapsis is through control of Mad2 and that Mad2 can bind other factors at the nuclear periphery. We also tested whether Mad2’s ability to adopt a specific conformation associated with its activity during spindle checkpoint function is required for its role in meiosis. A mutation that prevents Mad2 from adopting its active conformer fails to localize to the nuclear periphery, abolishes the synapsis checkpoint and exhibits substantial defects in meiotic synapsis. Thus, Mad2, and its regulation by Mad1, is a major regulator of meiotic synapsis in *C. elegans*.

**AUTHOR SUMMARY:** Sexual reproduction relies on production of gametes, such as eggs and sperm, which are produced during meiosis. During this specialized cell division, chromosomes replicate, pair with their homologs, undergo synapsis and finally undergo recombination, all of which are required for correct meiotic chromosome segregation. Chromosomes are highly mobile during these steps in meiosis but the specific role of this mobility is unclear. Here, we show that spindle assembly checkpoint proteins, Mad1 and Bub3, that regulate and monitor meiotic synapsis are implicated in chromosome movement, solidifying the functional link between chromosome mobility and synapsis. Moreover, we provide additional data that another spindle checkpoint effector, Mad2, and its regulation by Mad1, plays an important role in regulating meiotic synapsis.

## INTRODUCTION

Meiosis is a specialized biological process during which cells undergo a single round of DNA replication followed by two successive rounds of cell division. This process produces haploid gametes from diploid organisms. Diploidy is restored during sexual reproduction by the fusion of gametes, such as eggs and sperm, during fertilization, producing embryos. If chromosomes missegregate during meiosis, gametes and, upon their fertilization, embryos, will have the wrong number of chromosomes, also called aneuploidy. Aneuploidy during meiosis is frequently associated with miscarriages, infertility, and birth defects such as Down syndrome.

To ensure that chromosome segregation occurs normally during meiosis, critical events in meiotic prophase are tightly coordinated, monitored and regulated. Briefly, after replication, chromosomes pair with their homologs. Homologous interactions are stabilized by the assembly of a proteinaceous structure, the synaptonemal complex (SC) during a process called synapsis. Synapsis is a prerequisite for crossover recombination to generate linkages, or chiasmata, between homologs. These events are essential to direct proper meiotic chromosome segregation in which homologs and sister chromatids are separated during meiosis I and meiosis II respectively.

Because of their importance, multiple cell cycle checkpoints ensure the normal progression of synapsis and recombination, delay the cell cycle to correct errors and promote the removal of persistent abnormal cells (MacQueen and Hochwagen, 2011). One such checkpoint response, the synapsis checkpoint, triggers apoptosis to eliminate nuclei with unsynapsed chromosomes in *Caenorhabditis elegans* (Bhalla and Dernburg, 2005). This checkpoint relies on Pairing Centers (PCs), cis acting sites at one end of each chromosome promote pairing and synapsis (Bhalla and Dernburg, 2005; MacQueen et al., 2005). PCs play an important role, anchoring chromosome ends at the nuclear envelope to enable interaction with the SUN-1/ZYG-12 complex that spans the nuclear envelope; this interaction enables PCs to access the microtubule network in the cytoplasm (Labrador et al., 2013; Penkner et al., 2007; Sato et al., 2009), allowing chromosomes to become mobile within the nucleus. This mobilization is a conserved feature of meiotic prophase and essential for pairing and synapsis (Bhalla and Dernburg, 2008). Whether this mobilization also contributes to checkpoint function is unknown.

We recently showed that mitotic spindle assembly checkpoint (SAC) components Mad1, Mad2 and Bub3 are required to negatively regulate synapsis and promote the synapsis checkpoint response in *C. elegans* (Bohr et al., 2015). The genes that encode the *C. elegans* orthologs of Mad1 and Mad2 are *mdf-1* and *mdf-2*, respectively. However, for the sake of clarity, we will refer to these genes as *mad-1* and *mad-2* and their respective proteins as MAD-1 and MAD-2, consistent with *C. elegans* nomenclature. MAD-1 and MAD-2 localize to the nuclear envelope and interact with SUN-1, leading us to propose that these proteins may regulate and monitor synapsis through the ability of PCs to interact with and move at the nuclear envelope (Bohr et al., 2015). Indeed, PCs exhibit stereotypical behavior, called processive chromosome motions (PCMs), in which PCs travel continuously in a single direction for several seconds, often stretching chromosomes (Wynne et al., 2012). PCMs are dispensable for homolog pairing, reduce in frequency upon homolog synapsis and depend on the microtubule motor, dynein (Wynne et al., 2012). Since dynein is required for synapsis (Sato et al., 2009), PCMs have been suggested to trigger synapsis between accurate homolog pairs (Wynne et al., 2012).

Here we test whether spindle checkpoint components implicated in the synapsis checkpoint also affect PC movement, providing a potential link between chromosome mobility and the regulation and monitoring of meiotic synapsis. We find that SAC checkpoint mutants reduce the frequency of PCMs during meiotic prophase, consistent with the acceleration of synapsis observed in these mutant backgrounds. Further we investigate what functional aspects of SAC components are required for an efficient synapsis checkpoint. We show the N-terminal portion of MAD-1, required for the localization of the protein to the nuclear periphery, is also required for the synapsis checkpoint. However, unlike other mutant alleles of MAD-1, this inability to localize to the nuclear envelope does not affect MAD-2 localization or synapsis. In contrast, a mutation that affects MAD-1’s interaction with MAD-2 is crucial for MAD-2’s localization at the nuclear envelope, timely synapsis and a functional checkpoint. Finally, we demonstrate that the closed conformation of MAD-2 is required to regulate and monitor synapsis. Thus, MAD-2, and its regulation by MAD-1, seems to be a major regulator of meiotic synapsis in *C. elegans*.

## RESULTS

### PC movements are affected in SAC mutants

At the onset of meiosis, chromosomes are anchored to the nuclear envelope through their PCs. Prior to the entry to meiosis, the trans-membrane protein complex SUN-1/ZYG-12 is evenly distributed all around the nuclear envelope: SUN-1 faces the nucleus and mediates interaction with PCs and ZYG-12 faces the cytoplasm to mediate interactions with microtubules (Labrador et al., 2013; Penkner et al., 2007; Sato et al., 2009). The attachment of PCs to the SUN-1/ZYG-12 trans-membrane protein complex at the nuclear envelope leads to aggregation of the complex, which can be visualized cytologically as the formation of patches at the nuclear periphery (Baudrimont et al., 2010; Harper et al., 2011; Penkner et al., 2007; Sato et al., 2009). These patches are highly mobile, depend on microtubules for their mobility (Sato et al., 2009) and adopt two distinct modes of displacement: 1) periods when these patches are moving in different directions, while remaining close to their point of origin; and 2) processive chromosome motions (PCMs), where patches are continuously moving in the same direction for up to several seconds (Wynne et al., 2012). PCMs are not required for homolog pairing and reduce in frequency with synapsis, consistent with a role in regulating synapsis (Wynne et al., 2012). PCMs have been shown to occur approximately 15% of the time that SUN-1/ZYG-12 patches are visible (Wynne et al., 2012). These patches of SUN-1/ZYG-12 persist until synapsis is complete. After synapsis, the SUN-1/ZYG-12 complex is redistributed throughout nuclear envelope (Baudrimont et al., 2010).

In a previous work, we showed that MAD-1 and BUB-3 negatively regulate synapsis in a PC-dependent manner (Bohr et al., 2015). Given that PCMs correlate with the onset of synapsis and that chromosome mobility ceases when SC components are loaded prematurely on chromosomes (Zhang et al., 2012), we reasoned that MAD-1 and BUB-3’s inhibition of synapsis may be through regulation of PCMs. Therefore, we monitored PC movement in *mad-1* and *bub-3* mutants. We introduced the SUN-1-mRuby fusion protein (Rog and Dernburg, 2015) into *mad-1* and *bub-3* mutants and visualized PC movement using the two-dimensional assay previously developed (Wynne et al., 2012).

For all genotypes, we analyzed 3 to 5 nuclei from 2 to 3 wildtype germlines. In control animals, we detected 4-6 SUN-1-mRuby patches per nucleus with an average size of 0.65µm (Figure 1A), consistent with what has been previously reported (Wynne et al., 2012). When we analyzed SUN-1-mRuby patches in both *mad-1* and *bub-3* mutants, the number and size of patches were not different than that observed in control animals (Figure 1A), indicating that chromosome attachment did not appear perturbed.

**Figure 1:**
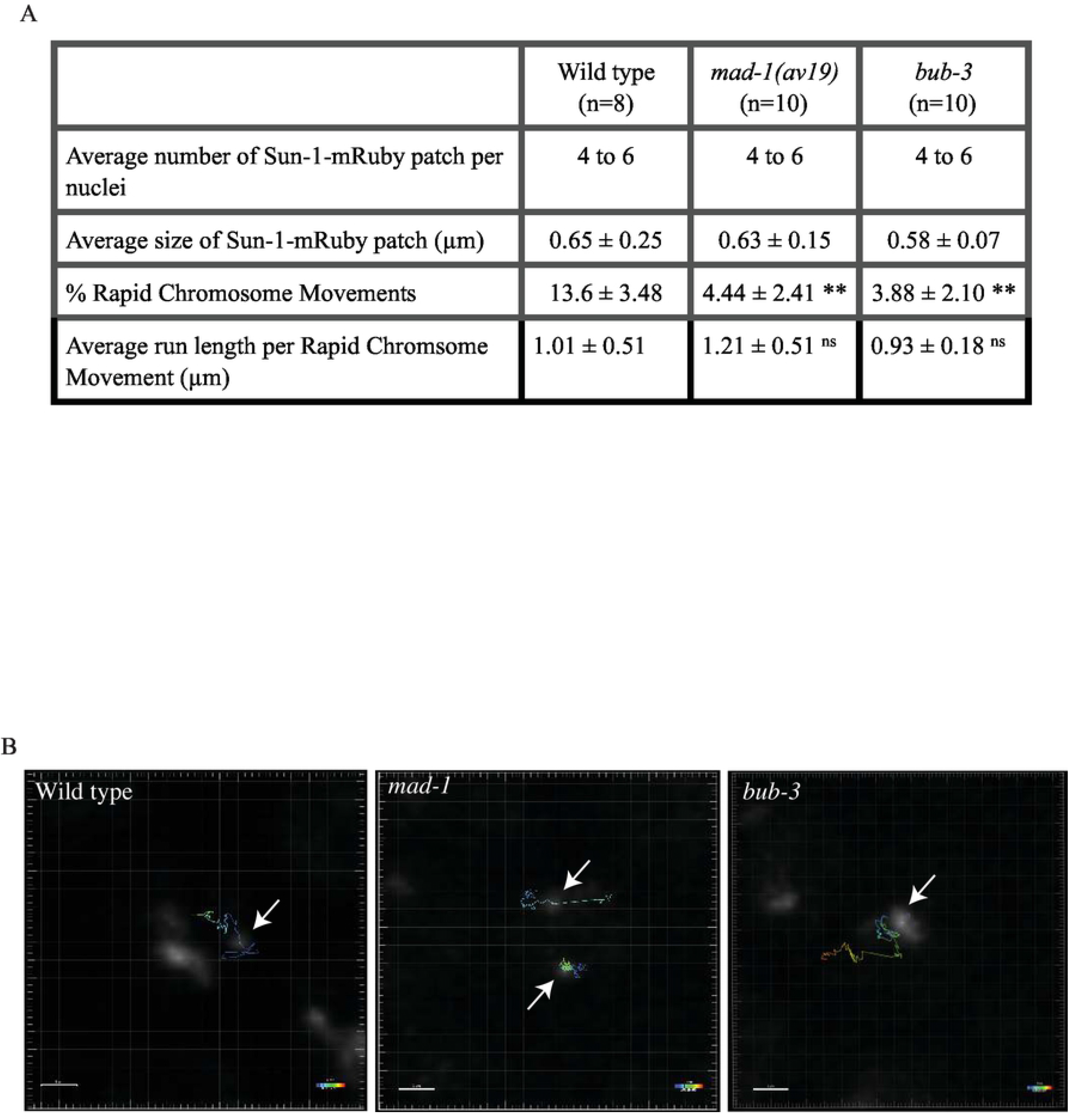
Spindle assembly checkpoint mutants affects progressive chromosome movements. A. Processive chromosome movements occur at lower frequency in *mad-1(av19)* and *bub-3* mutants. A ** indicates p value < 0.01 and ns indicates not significant. n indicated the number of nuclei analyzed. B. Images of SUN-1-mRuby patches and associated tracks of patch movement in wild type, *mad-1* and *bub-3* mutants. Arrows indicate tracked patches of SUN-1-mRuby. Bar: 1 µm.

We then wanted to analyze the mobility of SUN-1-mRuby patches, specifically PCMs. To analyze PCMs, we used criteria as defined in Wynne et al., 2012, in which SUN-1-mRuby patches are undergoing PCMs when they reach the minimum speed of 0.4µm/s for at least 1.2 sec. We found that that SUN-1-mRuby patches participated in PCMs (Video 1) for 13.6% of the time that they were mobile as patches in control worms, similar to the 15% observed (Wynne et al., 2012). The remaining time, patches exhibit short-range motions that are restricted to a small area. However, in contrast to control animals, we found that PCMs occur in *mad-1* (Video 2) and *bub-3* (Video 3) mutants but are only 4.44% and 3.88% of the observed PC movements respectively (Figure 1A). Despite being reduced in frequency, SUN-1-mRuby patches in *mad-1* and *bub-3* mutants traveled the same average distance when undergoing rapid chromosome movements as control animals (Figure 1A and B). Therefore, in these mutants, PCs still undergo stereotypical PCMs but at a reduced frequency, consistent with the accelerated synapsis we observe in *mad-1* and *bub-3* mutants.

### MAD-1’s localization to nuclear envelope is required for the synapsis checkpoint but not to regulate synapsis

Having established that PC movements are affected when some SAC components are mutated or deleted (Figure 1), we investigated which specific functions of some SAC components are necessary for the synapsis checkpoint and regulating synapsis. We previously showed that MAD-1 localizes to the nuclear periphery during meiotic prophase (Bohr et al., 2015),. Therefore, we tested whether this localization was required for monitoring and regulating synapsis (ΔN-MAD-1 in Figure S1). Amino acids 151 to 320 are required for MAD-1’s interaction with the nuclear pore component Tpr (NPP-21 in *C. elegans*) and its localization to the nuclear periphery in mitotic germline cells (Lara-Gonzalez et al., 2019). Deletion of this region also abrogated localization of MAD-1 at the nuclear periphery of meiotic germline nuclei (Figure 2A). In contrast to control animals with wildtype MAD-1, ΔN-MAD-1 adopted a diffuse localization inside nuclei and occupied area devoid of DNA (Figure 2A).

**Figure 2:**
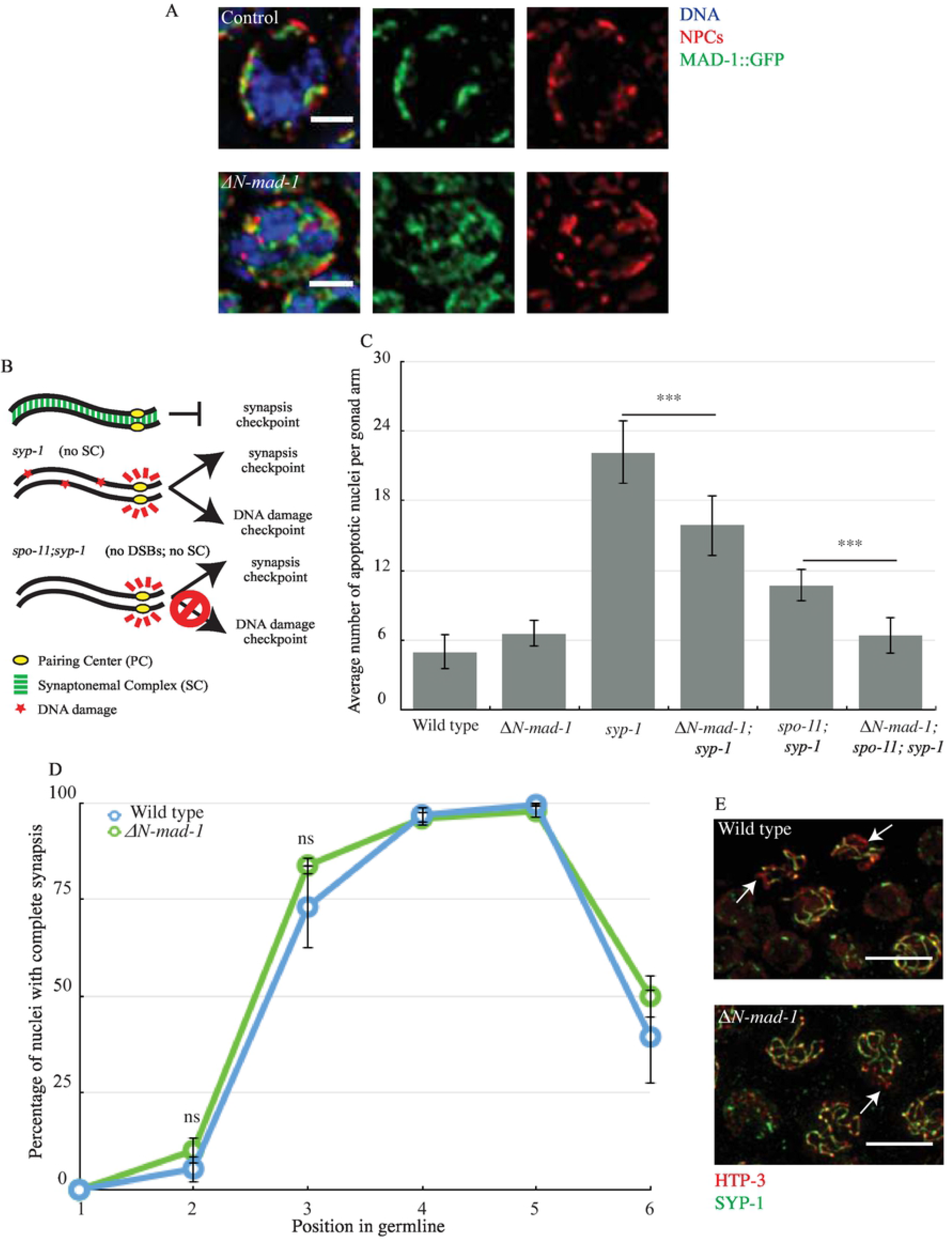
MAD-1’s localization to nuclear envelope is required for the synapsis checkpoint but not to regulate synapsis. A. ΔN-MAD-1 (green) localizes diffusely in the cytoplasm of meiotic nuclei and does not co-localize with NPCs (red). Images are partial projections of meiotic nuclei stained to visualize DNA (blue). Bar: 2 µm. B. A cartoon of meiotic checkpoints in *C. elegans*. C. *ΔN-mad-1* reduces germline apoptosis in *syp-1* and *spo-11; syp-1* mutants. A *** indicates p value < 0.0001. D. synapsis is unaffected in *ΔN-mad-1* mutants. ns indicates not significant. E. Images of nuclei during synapsis initiation in wild-type worms and *ΔN-ter-mad-1* mutants stained to visualize SYP-1 and HTP-3. Arrows indicates unsynapsed chromosomes. Bar: 5 µm.

Next, we tested what effect this deletion had on the synapsis checkpoint (Bohr et al., 2015). Synapsis is characterized by the assembly of a protein structure called synaptonemal complex (SC) between homologous chromosomes (Bhalla and Dernburg, 2008). In *C. elegans*, this protein structure is composed of a family of proteins, one of which is SYP-1. *syp-1* mutants do not load SC between homologues, producing unsynapsed chromosomes (MacQueen et al., 2002). In response to this abnormality, both the synapsis and DNA damage checkpoints are activated, resulting in very high levels of germline apoptosis (Figure 2B and C) (Bhalla and Dernburg, 2005). When we introduced the Δ*N-mad-1* deletion into the *syp-1* mutant background, the double mutant exhibited an intermediate level of germline apoptosis, indicating that the ability of MAD-1 to interact with Tpr and localize to the nuclear periphery is required for either synapsis or DNA damage checkpoint (Figure 2C). To determine which checkpoint is affected by the loss of the N terminus of MAD-1, we abolished the DNA damage checkpoint by using the *spo-11;syp-1* mutant background. SPO-11 generates double-strand breaks to initiate meiotic recombination (Dernburg et al., 1998); therefore, in this background only the synapsis checkpoint is activated (Figure 2B) (Bhalla and Dernburg, 2005). When we generate the Δ*N-mad-1;spo-11;syp-1* triple mutants, we observe wild-type levels of apoptosis, indicating the N terminus of MAD-1 is required for the synapsis checkpoint (Figure 2C).

We previously showed that in some spindle checkpoint mutants, a role in the synapsis checkpoint is coupled to a role in regulating synapsis. To determine whether this is also true for Δ*N-mad-1* deletion mutants, we assessed synapsis progression by staining for two SC proteins. We stained for HTP-3, an axial element that is loaded between sister chromatids before synapsis (MacQueen et al., 2005) and for SYP-1 (MacQueen et al., 2002). When we overlay HTP-3 and SYP-1 staining signals, stretches of HTP-3 without SYP-1 indicates the presence of unsynapsed chromosomes (arrows in Figure 2E) while colocalization of HTP-3 and SYP-1 indicates complete synapsis (Figure 2E). In *C. elegans*, meiotic nuclei in the germline are organized in a spatiotemporal gradient. Therefore, we divided germlines into six equivalent zones and calculated the percentage per zone of nuclei exhibiting complete synapsis to assay the progression of synapsis (Figure 2D). When we performed this analysis, Δ*N-mad-1* deletion mutants resembled wildtype germlines (Figure 2D), demonstrating that while the localization of MAD-1 to the nuclear envelope is required to monitor synapsis (Figure 2C), it is not required to regulate synapsis (Figure 2D). This is in contrast to other mutations in *mad-1* that both regulate and monitor synapsis (Bohr et al., 2015).

### MAD-1 is not required for MAD-2’s localization to the nuclear envelope in meiotic germline nuclei

Since Δ*N-mad-1* mutants did not affect synapsis (Figure 2D), unlike other *mad-1* mutants we had characterized (Bohr et al., 2015), we tested whether Δ*N-mad-1* deletion mutants affect the localization of another protein required for the synapsis checkpoint, MAD-2. MAD-2 adopts the same localization as MAD-1 in meiotic germline nuclei: the protein is targeted to the nuclear periphery in a punctate pattern (Bohr et al., 2015). We performed immunostaining using antibodies against nuclear pore complexes (NPCs) and MAD-2 and observed that, in contrast to a mutation that abolishes MAD-1’s checkpoint function (*mad-1[av19]*) and a null mutation in MAD-1 (*mad-1[gk2])*), MAD-2 localization to the nuclear periphery was unaffected by MAD-1’s absence from the nuclear periphery in Δ*N-mad-1* deletion mutants (Figure 3). As a control, we performed immunofluorescence against MAD-2 in *mad-2* null mutants (Figure 3). We also verified that MAD-2 localization was unaffected in *bub-3* mutants (Figure 3), having established a role for this gene in regulating and monitoring synapsis (Bohr et al., 2015) and affecting PCMs (Figure 1).

**Figure 3:**
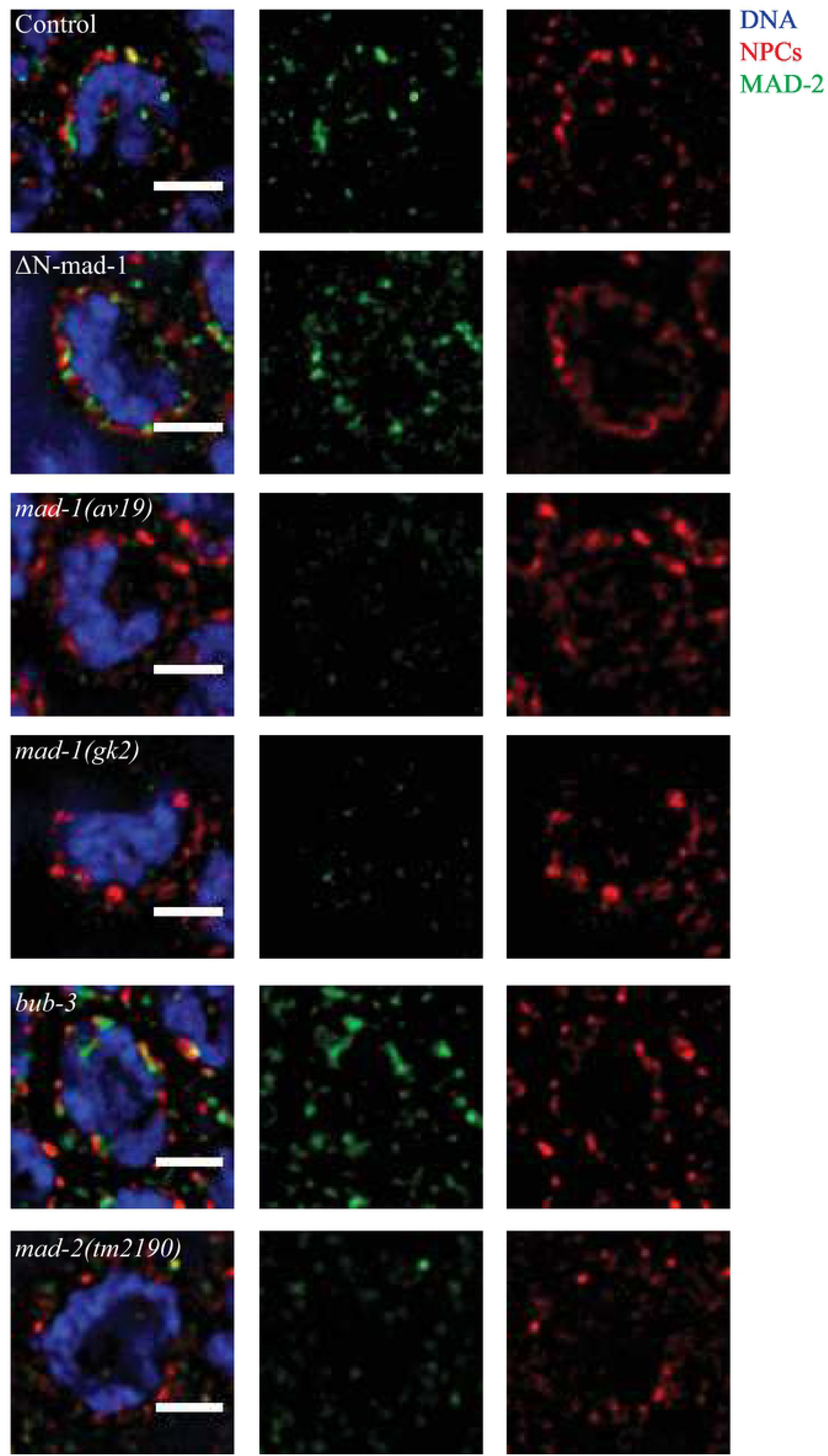
MAD-1 is not required for MAD-2’s localization to the nuclear envelop in meiotic germline nuclei. MAD-2 (green) co-localizes with NPCs (red) in *ΔN-mad-1* and *bub-3* mutants but is not detected in *mad-1(av19), mad-1(gk2)* and *mad-2(tm2190)* mutants. Images are partial projections of meiotic nuclei stained to visualize DNA (blue). Bar: 2 µm.

### MAD-1’s interaction with BUB-1 is not required to monitor or regulate synapsis

We had previously hypothesized that MAD-1’s localization to the nuclear periphery in meiotic germline nuclei suggested an interaction with PCs, cis-acting chromosomal regions essential for pairing, synapsis and synapsis checkpoint function (Bohr et al., 2015). In this way, we compared unsynapsed PCs to unattached kinetochores, which recruit Mad1 and Mad2 to initiate spindle assemble checkpoint signaling (Lara-Gonzalez et al., 2012). To further explore this connection, we took advantage of a mutation in Mad1 that prevents its localization to unattached kinetochores.

MAD-1 is recruited to unattached kinetochores through its interaction with BUB-1, a conserved kinase that is essential for chromosome segregation and spindle checkpoint function. We used a mutant version of MAD-1, *mad-1(E419A, R420A, D423A)* (Figure S1) that abolishes its binding to BUB-1, its localization to unattached kinetochores and its function in the spindle checkpoint (Moyle et al., 2014). We will refer to this allele as *mad-1(AAA)*. We tested if MAD-1’s ability to bind BUB-1 is also required for MAD-1 localization, checkpoint function and regulation of synapsis in meiosis. We stained fixed germlines with NPCs and MAD-1 antibodies and observed a localization comparable to wild type MAD-1 (Figure 4A). In this mutant, MAD-2 localization is also unaffected (Figure 4B). Therefore, the region of MAD-1 that is required to bind BUB-1 and localize to unattached kinetochores is not required for its localization to the nuclear periphery.

**Figure 4:**
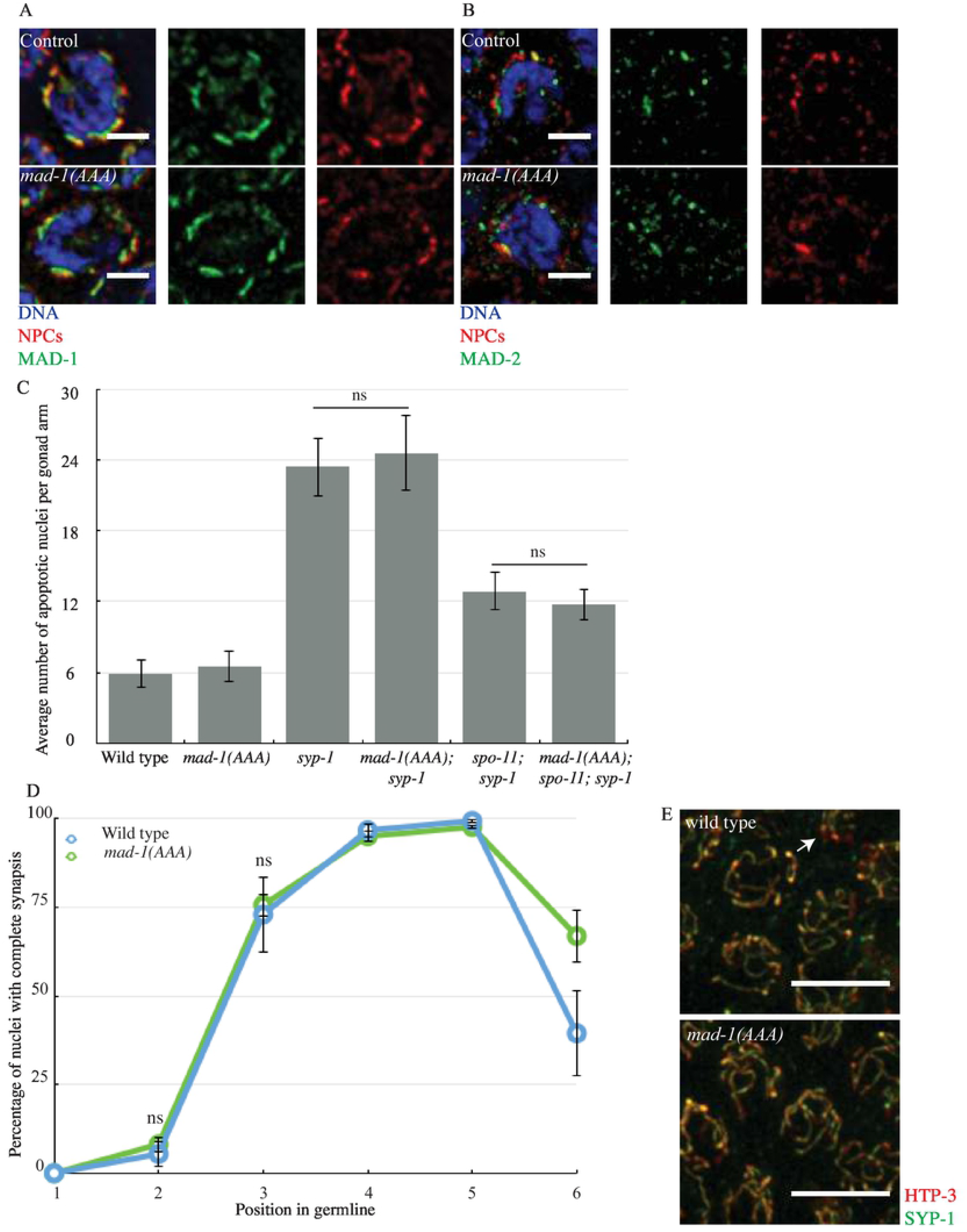
MAD-1’s interaction with BUB-1 is not required to monitor or regulate synapsis. A. MAD-1 (green) co-localizes with NPCs (red) in *mad-1(AAA)* mutants. Images are partial projections of meiotic nuclei stained to visualize DNA (blue). Bar: 2 µm. B. MAD-2 (green) co-localizes with NPCs (red) in *mad-1(AAA)* mutants. Images are partial projections of meiotic nuclei stained to visualize DNA (blue). Bar: 2 µm. C. The synapsis checkpoint and the DNA damage checkpoint are unperturbed in *mad-1(AAA)* mutants. D. synapsis is unaffected in *mad-1(AAA)* mutants. ns indicates not significant. E. Images of nuclei during synapsis initiation in wild-type and *mad-1(AAA)* mutants stained to visualize SYP-1 and HTP-3. Arrow indicates unsynapsed chromosomes.Bar: 5 µm.

Next, we tested whether this motif was required to regulate and monitors synapsis. We generated the double and triple mutants *syp-1;mad-1(AAA)* and *spo-11;syp-1;mad-1(AAA)*. When we assayed apoptosis, *syp-1 mad-1(AAA)* mutants were indistinguishable from *syp-1* single mutants. Similarly, *spo-11;syp-1;mad-1(AAA)* mutants were indistinguishable from *spo-11;syp-1* mutants (Figure 4C). This result indicates that neither the synapsis or DNA damage checkpoint are affected in the *mad-1(AAA)* mutants. When we assayed the progression of synapsis, synapsis in *mad-1(AAA)* mutants resembled synapsis in wildtype animals (Figure 4D). Thus, the motif required for MAD-1’s ability to interact with BUB-1 is not required for the synapsis checkpoint and does not regulate synapsis (Figure 4C and D).

### MAD-1’s ability to interact with MAD-2 is required to regulate and monitor synapsis

The correct localization of MAD-2 in Δ*N-mad-1* deletion mutants led us to consider the effects on regulating and monitoring synapsis if MAD-1 cannot bind MAD-2. We used a point mutation in *mad-1, mad-1(P504A)*, which abolishes its ability to bind MAD-2 (Figure S1) (Moyle et al., 2014). We will refer to this allele as *mad-1(A)* in this paper. First, we verified MAD-1’s localization in meiotic germline nuclei in this background. After staining for MAD-1 and NPCs, we were able to see that this point mutation does not affect the protein’s targeting to the nuclear periphery, similar to wildtype (Figure 5A) (Bohr et al., 2015; Stein et al., 2007; Yamamoto et al., 2008). Next, we looked at MAD-2 localization in this mutant background and were not able to detect the protein at the nuclear periphery (Figure 5B), similar to *mad-1(av19)* mutants and *mad-1(gk2)* null mutants (Figure 3). Thus, MAD-1’s ability to bind MAD-2 does not prevent MAD-1’s localization to the nuclear periphery in meiotic germline nuclei but does affect MAD-2’s.

**Figure 5:**
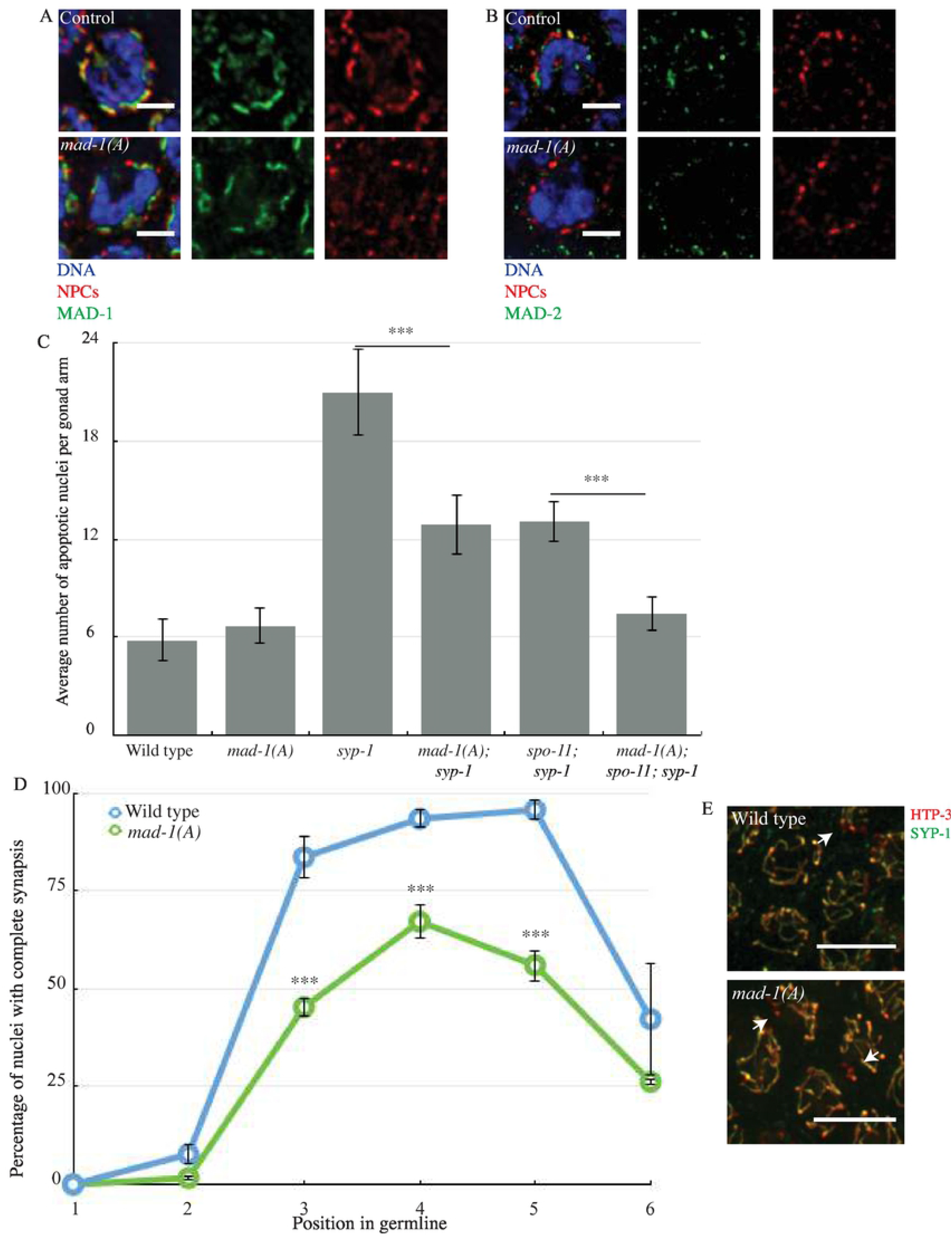
MAD-1’s ability to interact with MAD-2 is required to regulate and monitor synapsis. A. MAD-1(A) localizes at the nuclear periphery. B. MAD-2 (green) does not co-localize with NPCs (red) at the nuclear periphery in *mad-1(A)* mutants. Images are partial projections of meiotic nuclei stained to visualize DNA (blue). Bar: 2 µm. B. *mad-1(A)* reduces germline apoptosis in *syp-1* and *spo-11;syp-1* mutants. A *** indicates p value < 0.0001. C. Synapsis is reduced and delayed in *mad-1(A)* mutants. D. Images of nuclei during synapsis initiation in wild-type and *mad-1(A)* mutants stained to visualize SYP-1 and HTP-3. Arrows indicates unsynapsed chromosomes. Bar: 5 µm.

We then investigated the implication of MAD-1’s ability to bind MAD-2 for the synapsis checkpoint (Figure 5C). We combined *mad-1(A)* mutation in the *syp-1* background. We were able to observe an intermediate reduction in the number of apoptotic nuclei, indicating that one of the two checkpoints is affected by *mad-1(A)* mutation (Figure 5C). To determine which checkpoint is affected, we generated the triple mutant *mad-1(A); spo-11;syp-1*, which cannot activate the DNA damage checkpoint and only activates the synapsis checkpoint. Apoptosis was similar to wildtype in these triple mutants, indicating that MAD-1’s ability to bind MAD-2 is required for the synapsis checkpoint (Figure 5C).

Next, we investigated what effect this mutation had on synapsis. We observed that *mad-1(A)* mutants exhibit a dramatic delay of synapsis (Figure 5D, zones 2 and 3) and a reduction in the percentage of nuclei that complete synapsis (Figure 5D, zones 4 and 5, arrows in Figure 5E). Thus, MAD-1’s ability to bind MAD-2 is required to promote synapsis.

Since this role in promoting synapsis was unexpected, we were concerned that the synapsis defects we observed might be the indirect consequence of aneuploidy from defects in mitosis earlier in the germline. To test this, we attempted to detect aneuploidy in *mad-1(A)* mutant. We performed immunofluorescence with antibodies against HIM-8 to identify aneuploid nuclei that either had no HIM-8 staining or more than two HIM-8 foci (Figure S2). We did not observe any nuclei with no HIM-8 or more than two HIM-8 signals in this mutant background, arguing against defects in ploidy and supporting a role for MAD-1’s ability to bind MAD-2 in regulating timely synapsis.

To further address this possibility, we scored apoptosis in *mad-1(A)* single mutants. Defects in mitotic checkpoint function in mitotic germline nuclei can produce aneuploidy in meiotic nuclei that activate the DNA damage checkpoint and elevate apoptosis (Stevens et al., 2013). However, the level of apoptosis in *mad-1(A)* single mutants was comparable to wildtype animals (Figure 5C), supporting our hypothesis that the synapsis defects we observe in *mad-1(A)* mutant are not a consequence of defects in the mitotic region of the germline and are likely not severe enough to activate the DNA damage checkpoint, similar to other mutant backgrounds that exhibit asynapsis in a subset of meiotic nuclei (Bhalla and Dernburg, 2005; MacQueen et al., 2005). All together, these data indicate that MAD-1’s ability to interact with MAD-2 is important for MAD-2 localization to the nuclear periphery but not for MAD-1 targeting to the nuclear periphery. Further, this interaction is required to promote the synapsis checkpoint and synapsis. This is in contrast to *mad1* null and *mad-1(av19)* mutants, which promote the synapsis checkpoint but inhibit synapsis (Bohr et al., 2015).

### MAD-2’s ability to adopt the closed conformation is required to regulate and monitor synapsis

MAD-2 is essential for the spindle checkpoint and the synapsis checkpoint. Its role in the spindle checkpoint has been extensively characterized (Lara-Gonzalez et al., 2012). MAD-2 adopts two conformations, an open and a closed conformation, depending on whether it is binding other protein partners (Rosenberg and Corbett, 2015). The open version is unbound and inactive in the spindle checkpoint. MAD-2 adopts the closed version upon binding MAD-1 (Luo et al., 2004; Sironi et al., 2002) either at the nuclear envelope (Rodriguez-Bravo et al., 2014) or at unattached kinetochores (Chen et al., 1996, 1998; Li and Benezra, 1996; Sironi et al., 2001). It also adopts the closed conformation when bound to Cdc20 (De Antoni et al., 2005; DeAntoni et al., 2005; Luo et al., 2002), during formation of the mitotic checkpoint complex. Thus, the closed version of MAD-2 is the active conformer during spindle checkpoint function. Recent work has shown that when MAD-2 is mutated so that it cannot convert to the closed conformation and remains locked in its open conformation, this mutant version of the protein cannot support the spindle checkpoint and is no longer detected at unattached kinetochores (De Antoni et al., 2005; DeAntoni et al., 2005; Lara-Gonzalez et al., 2021; Nezi et al., 2006). To evaluate the importance of this conversion for its meiotic role, we used a *mad-2* mutant that is locked in the open conformation *(mad-2[V193N])*; we will refer to this allele as *mad-2-open*.

First, we determined how this mutation affected the protein’s localization. When we stained germlines with NPCs and MAD-2 antibodies in *mad-2-open* mutants, we could not detect the protein in meiotic nuclei (Figure 6A), indicating that MAD-2’s ability to adopt the closed conformer is required for its localization to the nuclear periphery.

**Figure 6:**
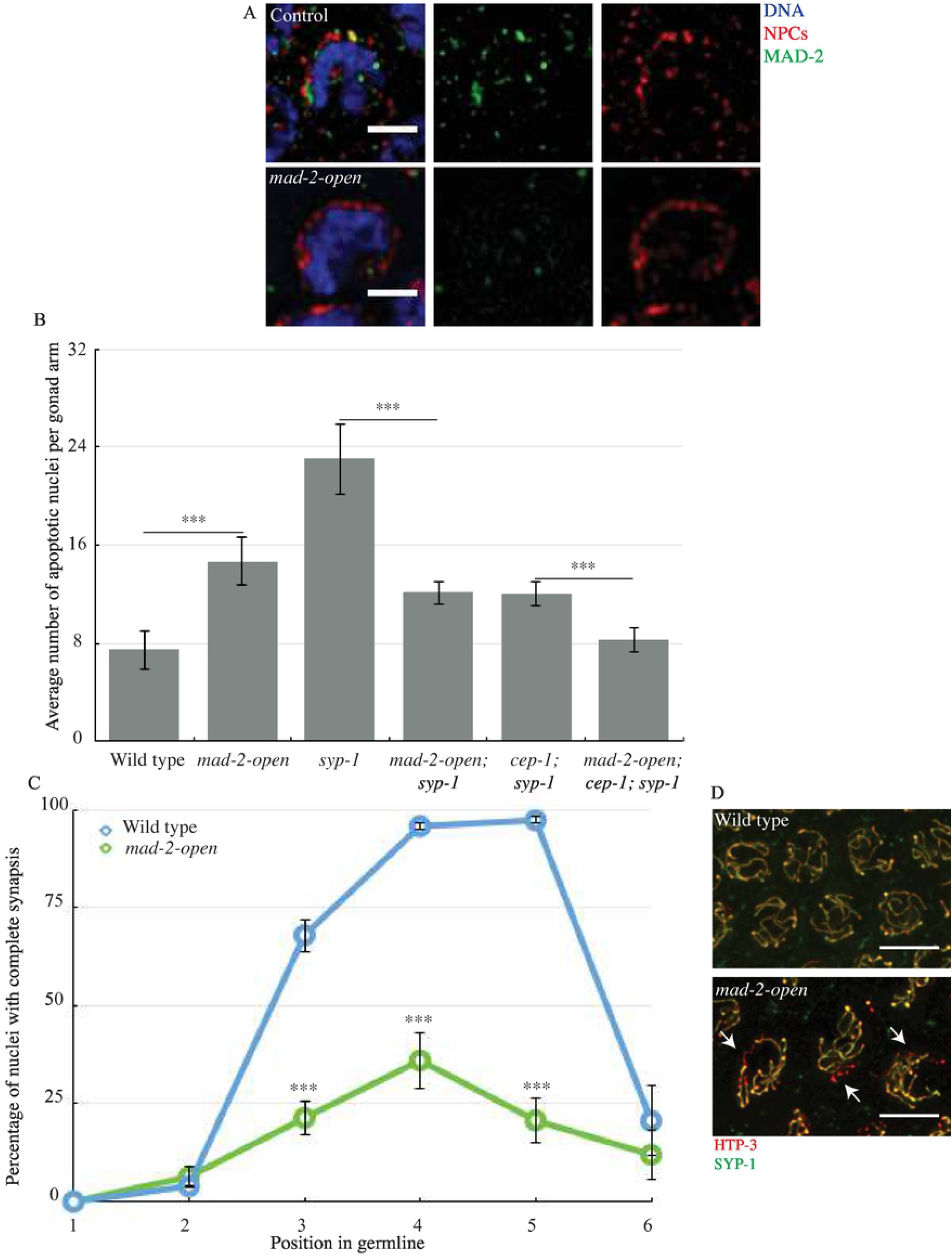
MAD-2’s ability to adopt the closed conformation is required to regulate and monitor synapsis. A. MAD-2 (green) does not co-localize with NPCs (red) at the nuclear periphery when the protein is locked in the open conformation. Images are partial projections of meiotic nuclei stained to visualize DNA (blue). Bar: 2 µm. B. *mad-2-open* reduces germline apoptosis in *syp-1* and *cep-1;syp-1* mutants. A *** indicates p value < 0.0001. C. Synapsis is reduced and delayed when Mad2 is locked in open conformation. A *** indicates p value < 0.0001. D. Images of nuclei during synapsis initiation in wild-type and *mad-2-open* mutants stained to visualize SYP-1 and HTP-3. Arrows indicates unsynapsed chromosomes. Bar: 5 µm.

Next, we evaluated its role in the synapsis checkpoint. We introduced this mutation into *syp-1* mutants and assayed apoptosis. When compared to the *syp-1* single mutant background, the *syp-1*;*mad-2-open* double mutants exhibit an intermediate level of apoptosis (Figure 6B), indicating that either the synapsis checkpoint or the DNA damage checkpoint is affected. Because the *mad-2* gene is closely linked to *spo-11*, we used *cep-1* to prevent DNA damage checkpoint-induced apoptosis in *mad-2-open* mutants. *cep-1* is the *C. elegans* ortholog of p53 and is required for the DNA damage response (Derry et al., 2001; Schumacher et al., 2001). We generated the *mad-2-open;cep-1;syp-1* triple mutant to clarify which checkpoint is affected. We observed a wild type level of apoptosis in *mad-2-open;cep-1;syp-1* triple mutants, indicating the ability to adopt the closed conformation is required for the synapsis checkpoint (Figure 6B).

Having established that this mutant disrupted the synapsis checkpoint, we assessed its effect on synapsis (Figure 6C and D). We observed a dramatic delay and reduction in the percentage of nuclei that completed synapsis in *mad-2 open* mutants. This phenotype is more severe than the one observed for *mad-1(A)* mutant (Figure 5D). In *mad-1(A)* mutants, 70% of meiotic nuclei complete synapsis in zone 4, while in *mad-2-open* mutants, only 40% do (Figure 5D and Figure 6C and D). Since complete synapsis is required for the proper progression of DNA repair and meiotic recombination, this severe defect in synapsis likely explains the elevated apoptosis we observe in *mad-2-open* single mutants (Figure 6B). Indeed, when *mad-2-open;cep-1* double mutants are generated and apoptosis assayed, the level of apoptosis is similar to *cep-1* single mutants and significantly lower than *mad-2-open* single mutants, indicating that *mad-2-open* mutants activate the DNA damage checkpoint (Figure S3).

Similar to our analysis of *mad-1(A)* mutants, we wondered if some of this asynapsis in *mad-2-open* mutants was the consequence of aneuploidy in meiotic nuclei. To assess this, we stained *mad-2-open* meiotic nuclei with antibodies against the X chromosome PC protein, HIM-8. We could detect nuclei that either contained no HIM-8 foci or more than two, indicating aneuploidy of the X chromosome (Figure S2). When we quantified this defect, we observed it in 3% of meiotic nuclei. Therefore, some small proportion of unsynapsed chromosomes in meiotic nuclei are likely the product of aneuploidy and not strictly a defect in synapsis in *mad-2-open* mutants. However, even if we assigned comparable rates of aneuploidy to the remaining five autosomes, this degree of aneuploidy is unlikely to explain the dramatic defect in synapsis that we observe *mad-2-open* mutants.

## DISCUSSION

When we first reported that spindle checkpoint proteins played a role in regulating and monitoring synapsis, we hypothesized that spindle checkpoint proteins might regulate the dynamics of PCs at the nuclear envelope, given the relationship between synapsis initiation and chromosome mobility (Bohr et al., 2015). Our analysis of chromosome movements in *mad-1* and *bub-3* mutants validates this hypothesis, demonstrating that stereotypical PC behaviors, namely processive chromosome movements (Wynne et al., 2012), show the same hallmark features in *mad-1* and *bub-3* mutants as wildtype animals but are reduced in frequency (Figure 1). Whether this reduction in PCM frequency is a cause or consequence of the accelerated synapsis we observe in these mutant backgrounds is still an open question.

The spindle checkpoint, and the functional requirements of its essential factors, has been studied extensively (Lara-Gonzalez et al., 2012). We took advantage of these studies to test what aspects of MAD-1 and MAD-2 function are required for the regulation and monitoring of synapsis. Somewhat surprisingly, we found that a mutation that abolished MAD-1’s association with the nuclear envelope (Lara-Gonzalez et al., 2019) did not affect MAD-2 localization (Figure 2A and Figure 3), indicating that MAD-2 can bind additional factors at the nuclear envelope besides MAD-1 during meiosis. MAD-2 has been shown to bind the insulin receptor and regulate its internalization dynamics in mice (Choi et al., 2016), raising the possibility that MAD-2 may bind other factors at the nuclear envelope in other developmental contexts as well. Further, despite the absence of

MAD-1, the presence of MAD-2 at the nuclear envelope still promotes the timely progression of synapsis (Figure 2D), suggesting that MAD-1’s primary role in regulating synapsis is through control of MAD-2.

This interpretation is borne out by our analysis of a *mad-1* mutant that no longer binds MAD-2, *mad-1(A*) (Moyle et al., 2014). This mutant protein is localized to the nuclear envelope (Figure 5A) but MAD-2 is not (Figure 5B), indicating that although MAD-1 may not be required for MAD-2’s localization to the nuclear envelope, this interaction is required for MAD-2’s presence inside the nucleus. This suggests a potential regulatory role for MAD-1 in shuttling MAD-2 into meiotic nuclei to carry out its role in regulating and monitoring synapsis.

We were surprised to observe that *mad-1(A)* mutants, unlike *mad-1* null or *mad-1(av19)* mutants, delay synapsis (Figure 5D). We ruled out the possibility that this was a consequence of the spindle checkpoint defect resulting in aneuploidy in meiotic cells (Figure S2). Further, since *mad-1(AAA)* mutants also have a spindle checkpoint defect (Moyle et al., 2014) and don’t affect synapsis (Figure 4D), we are comfortable attributing these phenotypes to a meiotic defect. These data suggest when MAD-2 cannot bind MAD-1, MAD-2 acts as a gain of function, disrupting synapsis. We speculate that this unregulated population of MAD-2 is now competent to bind additional meiotic factors, such as CMT-1 and/or PCH-2 (Deshong et al., 2014; Giacopazzi et al., 2020) that it is normally prevented from interacting with during meiosis. Indeed, the amount of non-homologous synapsis we observe in *mad-1(A)* mutants, ∼4%, is similar to what is observed in *cmt-1* null mutants (Giacopazzi et al., 2020), consistent with this possibility. Given that MAD-2 interacts with these factors during mitotic spindle checkpoint function (Nelson et al., 2015), MAD-2’s sequestration during meiosis may be an important regulatory event to promote meiotic synapsis.

Finally, we’ve shown that MAD-2’s ability to adopt its closed conformation is important for its localization to the nuclear envelope (Figure 6A), its role in the synapsis checkpoint (Figure 6B) and its regulation of synapsis (Figure 6C). One of the proteins it complexes with to adopt its closed conformation is definitely MAD-1, as demonstrated by MAD-2 absence from the nuclear envelope in *mad-1(A)* mutants (Figure 5B). However, MAD-2’s continued presence at the nuclear envelope in Δ*N-mad-1* mutants (Figure 3) illustrates that MAD-2 complexes with some other factor at the nuclear envelope during meiotic prophase and this has important implications for the regulation and monitoring of synapsis in *C. elegans*. Identifying this factor is an important next step in understanding MAD-2’s meiotic function.

Despite the effect of spindle checkpoint mutants on PC movement (Figure 1) and our previous model that spindle checkpoint mutants regulate and monitors meiotic synapsis by assessing whether PCs at the nuclear envelope are synapsed (Bohr et al., 2015), it’s unlikely that the role of spindle checkpoint proteins in regulating and monitoring meiotic synapsis at unsynapsed PCs can be compared with their role at unattached kinetochores. First, while a mutation that prevents MAD-1’s localization to the nuclear envelope, *ΔN-mad-1*, abrogates the synapsis checkpoint (Figure 2C), it does not affect synapsis (Figure 2D), indicating that MAD-1’s absence from the nuclear envelope does not affect the progression of synapsis. Further, the uncoupling of the regulation and monitoring of synapsis in Δ*N-mad-1* mutants indicates that its role in the checkpoint does not depend on its localization to the nuclear envelope, in direct contrast to our model. It is formally possible that MAD-1’s dispensability in regulating synapsis is because of MAD-2’s continued presence at the nuclear envelope in this mutant background (Figure 3). However, we do not favor this possibility based on MAD-2’s absence at the nuclear envelope and the dramatic defect in synapsis we observe in *mad-2-open* mutants (Figure 6A and C). If our model was correct, we might have predicted that *mad-2-open* mutants would accelerate synapsis, similar to *mad-1(av19)* and *mad-1* null mutants, which also fail to localize MAD-2 at the nuclear envelope (Figure 3). Instead, these data suggest a more complicated role for spindle checkpoint proteins in regulating and monitoring synapsis than we had previously proposed. Understanding this role may further expand the repertoire of spindle checkpoint proteins beyond their well-characterized roles in regulating the cell cycle and monitoring kinetochore attachment.

## MATERIALS AND METHODS

### Genetics and worm strains

The wild type *C. elegans* strain background was Bristol N2 (Brenner, 1974). All experiments were performed on adult hermaphrodites at 20°C under standard conditions unless otherwise stated. Mutations and rearrangements used were as follows:

LG I: *cep-1(gk138)*

LG II: *bub-3(ok3437), mln1 [mls14 dpy-10(e128)], ltSi609[pOD1584/pMM9; Pmdf-*

*1::mdf-1(P504A)::mdf-1 3*’*UTR; cb-unc-119(+)], ltSi620[pOD1595/pMM13; pmdf-*

*1::GFP::mdf1(E419A, R420A, D423A)::mdf1 3*’*UTR; cb-unc-119(+)], ltSi677*

*[pPLG034; Pmdf-1::GFP::mdf-1(Δ151–320)::mdf-1 3′UTR; cb-unc-119(+)]*,

*ltSi1514[pPLG333; Pmdf-2::mdf-2 delta hairpin intron 4 V193N::mdf-2 3’UTR; cb-unc-119(+)]*

LG III: *unc-119(ed3)*

LG IV: *mdf-2(tm2190), spo-11(ok79), nT1[unc-?(n754let-?(m435)]*

LG V: *mdf-1(av19), mdf-1(gk2), syp-1(me17), bcIs39[Plim-7::ced-1::gfp; lin-15(+)]*,

*ieSi21 [Psun-1::sun-1::mRuby::sun-1 3’UTR + Cbr-unc-119(+)], dpy-11(e224), nT1[unc-?(n754let-?(m435)]*

### Quantification of germline apoptosis

Scoring of germline apoptosis was performed as previously described in Bhalla and Dernburg, 2005. L4 hermaphrodites were allowed to age for 22 h at 20°C. Live worms were mounted under coverslips on 1.5% agarose pads containing 0.2 mM levamisole for wild type and moving strains or 0.1 mM levamisole for *dpy* strains. Minimum of 20 germlines were analyzed for each genotype by performing live fluorescence microscopy and counting the number of cells fully surrounded by CED-1::GFP. All experiments were performed three times. Significance was assessed using a paired *t*-test.

### Antibodies, immunostaining and microscopy

Immunostaining was performed on worms 20 to 24 f after L4 stage. Gonad dissection were performed in 1x EBT (250 mM Hepes-Cl, pH 7.4, 1.18 M NaCl, 480 mM KCl, 20 mM EDTA, 5 mM EGTA) + 0.1% Tween 20 and 20 mM sodium azide. An equal volume of 2% formaldehyde in EBT (final concentration was 1% formaldehyde) was added and allowed to incubate under coverslip for 5 min. The sample was mounted on HistoBond slides (75 × 25 × 1 mm from VWR), freeze-cracked, and immediately incubated in methanol at -20°C for 1 min and transferred to PBST (PBS with Tween20). After a total of 3 washes of PBST, the samples were incubated for 30 min in 1% bovine serum albumin diluted in PBST. A hand-cut paraffin square was used to cover the tissue with 50 µL of antibody solution. Incubation was conducted in a humid chamber at 4°C overnight. Slides were rinsed 3 times in PBST and incubated for 2 h at room temperature with fluorophore-conjugated secondary antibody solution at a dilution of 1:500. Samples were rinsed in PBST, DAPI stained in PBST (5 µg/mL) and rinsed a last time in PBST. Samples were then mounted in 12 µL of mounting media (20 M N-propyl gallate [Sigma-Aldrich] and 0.14 M Tris in glycerol) with a no. 1.5 (22 mm^2^) coverslip, and sealed with nail polish.

Primary antibodies were as follows (dilutions are indicated in parentheses). Rabbit anti-SYP-1 (1:500; MacQueen et al., 2002), chicken anti-HTP-3 (1:250; MacQueen et al., 2005), rabbit anti-MAD-2 and anti-MAD-1 (1:10000; Essex et al., 2009), mouse anti-NPC MAb414 (1:5000; Covance; Davis and Blobel, 1986), rat anti-HIM-8 (1:2500; Phillips and Derburg 2006) and goat anti-GFP (1:10000; Hua et al., 2009) Antibodies against SYP-1 were provided by A. Villeneuve (Stanford University, Palo Alto, CA). Antibodies against HTP-3 and HIM-8 were provided by A. Dernburg (University of California Berkley/E.O. Lawrence Berkley National Lab, Berkley, CA). Antibodies against MAD-1 and MAD-2 were provided by A. Desai (Ludwig Institute/University of California, San Diego, CA). Antibodies against GFP were provided by S. Strome (University of California, Santa Cruz, CA).

Secondary antibodies were Cy3, Cy5 and Alexa Fluor 488 anti-mouse, anti-rabbit, anti-guinea pig, anti-rat and anti-chicken (1:250; Jackson ImmunoResearch Laboratories, Inc.)

Quantification of synapsis was performed with a minimum of three whole germlines per genotype as in Phillips et al. (2005) on animals 24 h after L4 stage. The gonads were divided into six equal-sized regions, beginning at the distal tip of the gonad and progressing through the end or late pachytene.

All images were acquired at room temperature using a Delta-Vision Personnal DV system (GE Healthcare) equipped with a 100x NA 1.4 oil immersion objective (Olympus), resulting in an effective xy pixel spacing of 0.064 or 0.040 µm. Images were captured using a charge-coupled device camera (Cool-SNAP HQ; Photometrics). Three-dimensional images stacks were performed using functions in the softWoRx software package. Projections were calculated by a maximum intensity algorithm. Composite images were assembled, and some false coloring was performed with Fiji and Photoshop software (Adobe).

### Live imaging

For time lapse imaging of meiosis, we followed the protocol as described in Wynn et al., 2012. Briefly, young adult worms (16-20 h after L4 stage) were immobilized on freshly made 3% agarose pad in a drop of M9 media containing 0.4 mM (0.05%) tetramisole (Sigma Aldrich) and 3.8 mM (0.5%) tricaine (Sigma Aldrich). A 22 × 22 × 0.17 mm coverslip (Schott nexterion) was applied after 2 min of immersion in the anesthetic media. The monolayer of meiotic nuclei closest to the coverslip was imaged and collected at 20°C or room temperature no longer than 15 min after immersion. Images were acquired on a Solamere spinning disk confocal system piloted by µManager software (Edelstein et al., 2014) and equipped with a Yokogawa CSUX-1 scan head, a Nikon (Garden City, NY) TE2000-E inverted stand, a Hamamatsu ImageEM × 2 camera, LX/MAS 489 nm laser attenuated to 10%, and a Plan Apo × 60/1.4 numerical aperture oil objective.

For 2D confocal imaging, a focal plane near apical surface of many nuclei was imaged to 50-100 ms exposure at 489 nm with images acquisition every 400 ms for ≤80 s. These settings were used for rapid chromosome movements collection data.

For quantification and size measurement of SUN-1-mRuby patches, we imaged the first layer of nuclei in live worms. We exposed the germlines at 489 nm for 50-100 ms.

### Processive chromosome movements detection

We determined processive chromosome movements as when a SUN-1-mRuby patch moved in a continuous direction with a speed over 0.4 µm/sec for at least 3 consecutive time points (1.2 sec), as defined by (Wynne et al., 2012).

## ACKNOWLEDGEMENTS

We would like to thank Pablo Lara-Gonzalez, Arshad Desai, Karen Oegema, Abby Dernburg, and Anne Villeneuve for valuable strains and reagents. This work was supported by the NIH (grant number R01GM097144 [N.B.]). Some strains were provided by the CGC, which is funded by NIH Office of Research Infrastructure Programs (P40 OD010440).

## Author Contributions

Conceptualization and Methodology, A.D. and N.B.; Investigation, A.D.; Writing -Original Draft, A.D. and N.B.; Writing -Review & Editing, A.D. and N.B.; Supervision and Funding Acquisition, N.B.

## Declaration of Interests

The authors declare no competing interests.

## Figure Legends

**Figure S1:**
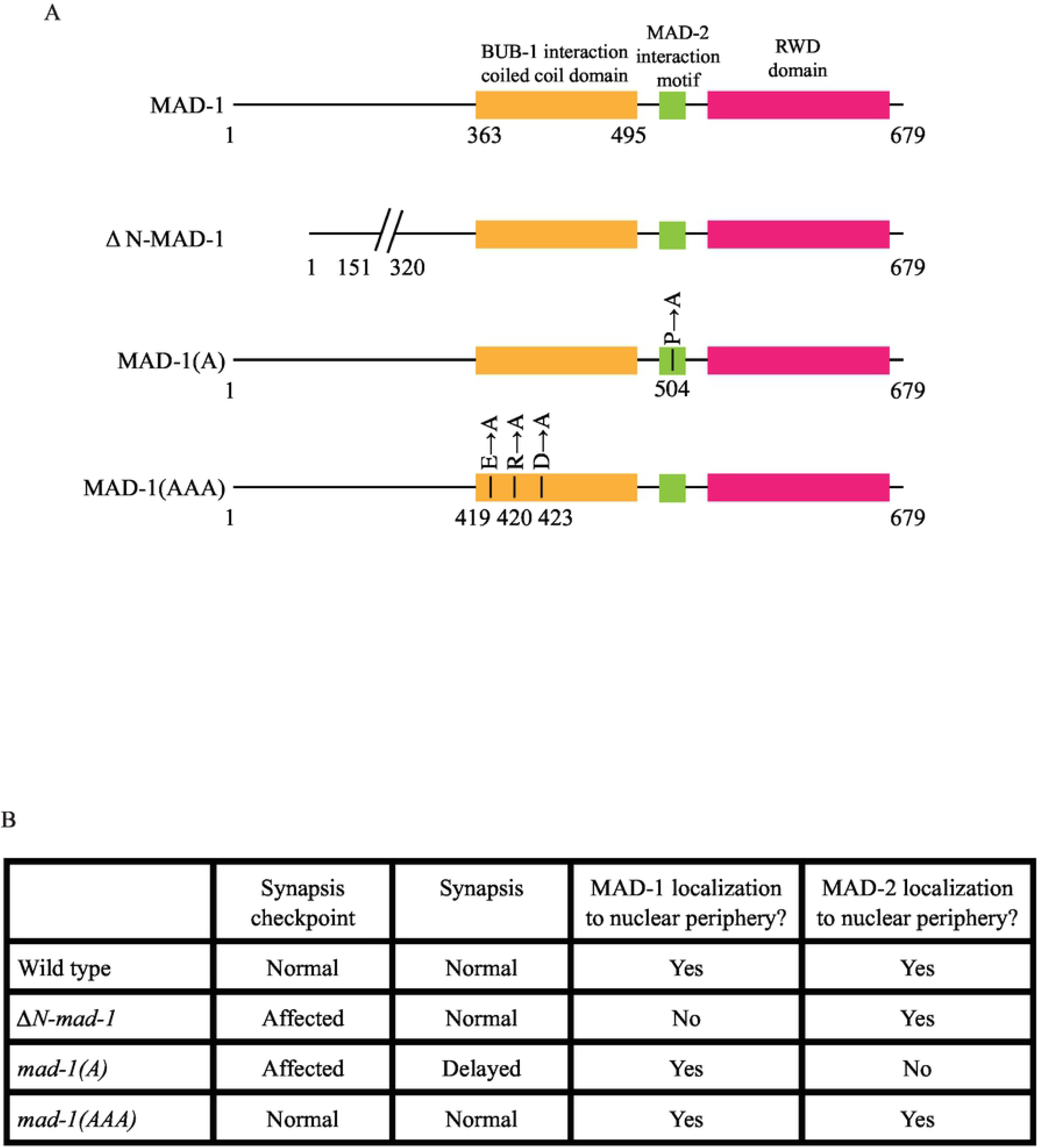
Summary of *mad-1* mutants studied in this paper. A. Cartoon of the different *mad-1* mutants studied in this paper. B. Summary of observed phenotypes.

**Figure S2:**
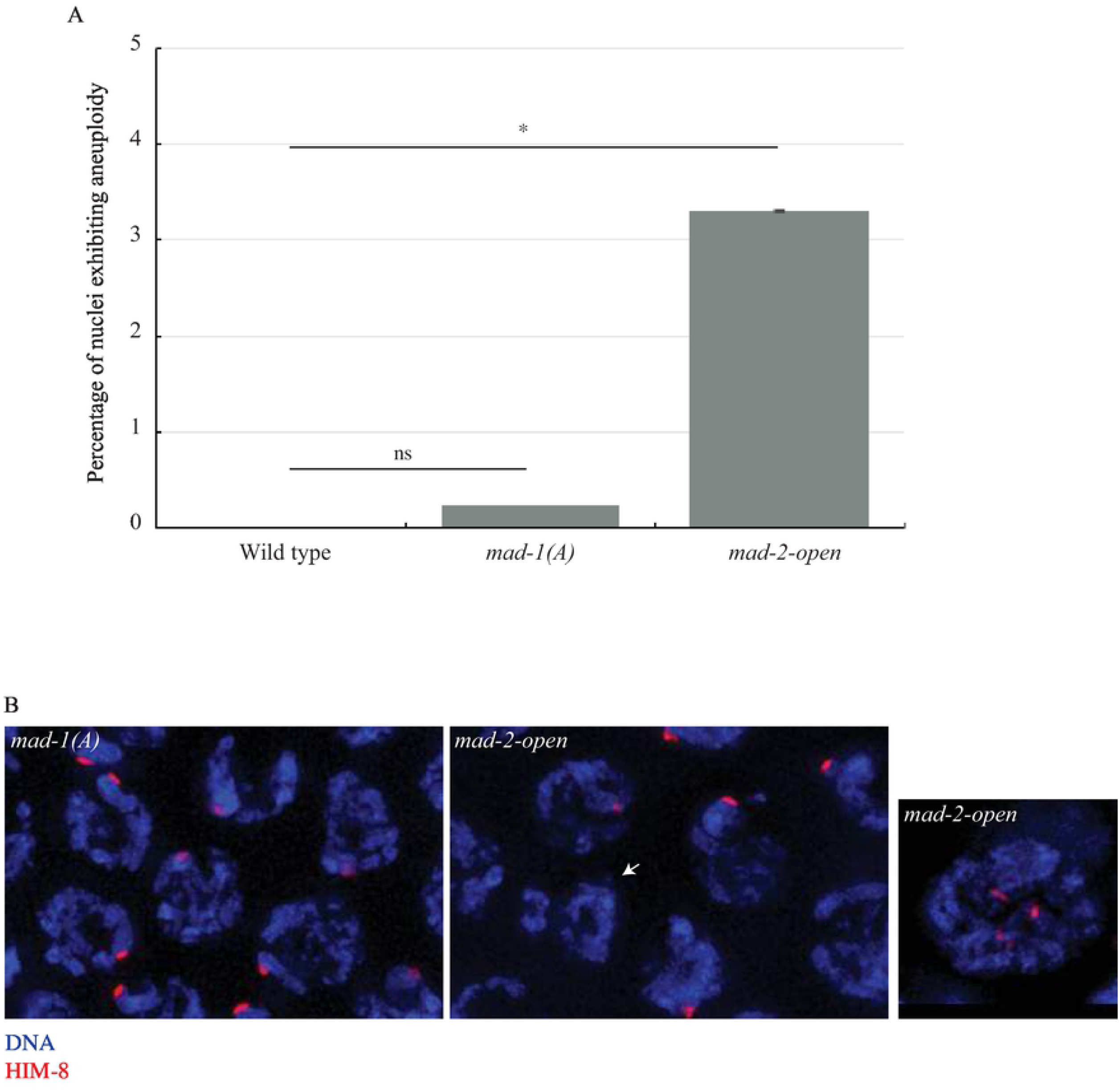
Aneuploidy is observed in *mad-2-open* but not *mad-1(A)* mutants. A. Meiotic nuclei exhibit aneuploidy in *mad-2-open* mutants but not in *mad-1(A)* mutants. B. Example of nuclei exhibiting aneuploidy in *mad-2-open* mutants. Images are projections of meiotic nuclei stained to visualize DNA (blue) and HIM-8 protein (red). Arrows indicates nuclei with no HIM-8 foci.

**Figure S3:**
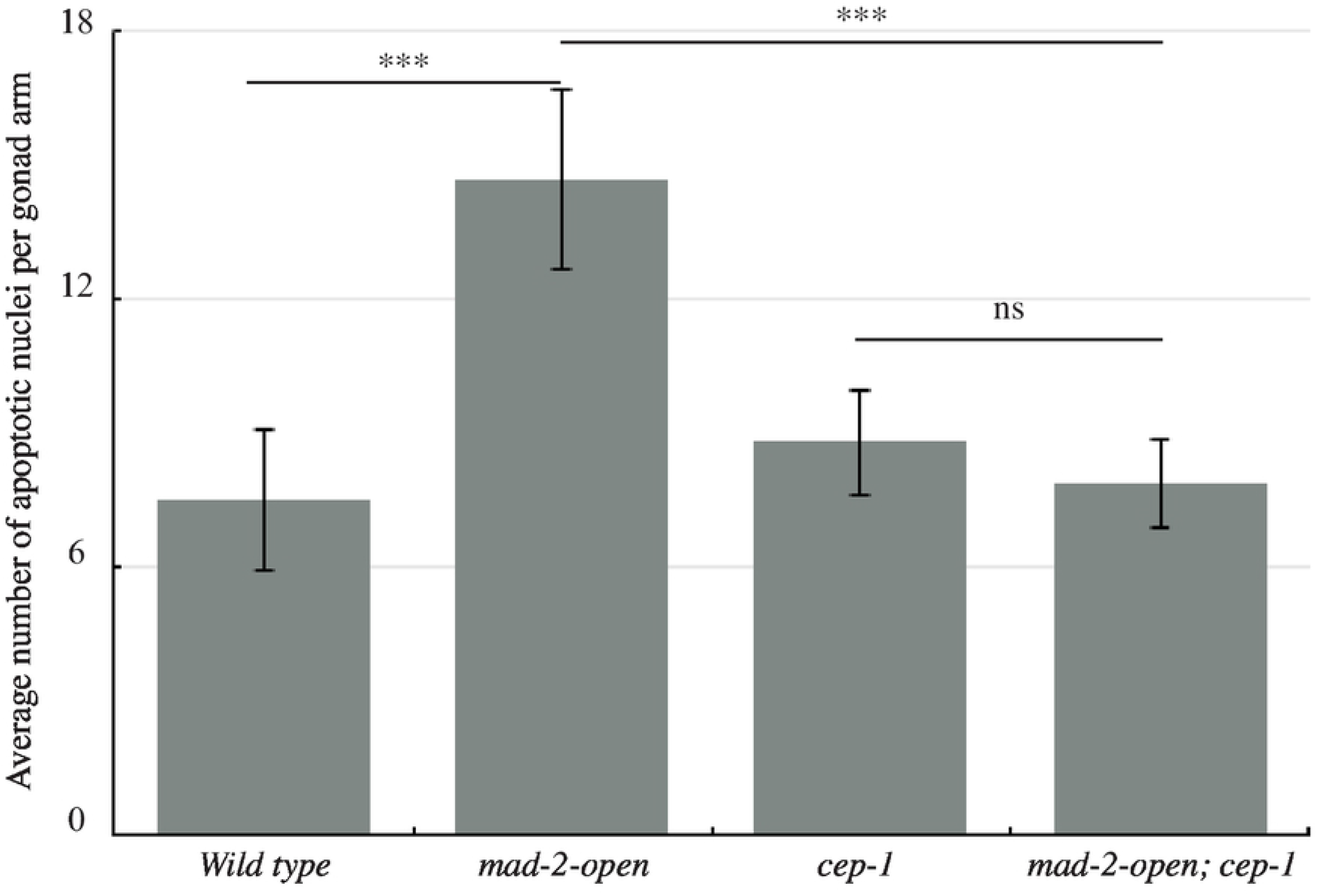
*mad-2-open* mutants activate the DNA damage checkpoint. *cep-1* reduces apoptosis in *mad-2-open* mutants. A *** indicates p value < 0.0001 and ns indicates not significant.

## Video Legends

**Video 1:** Movement of a SUN-1-mRuby patch and its trajectory in a control nucleus. Movie is displayed at 10 frames per second.

**Video 2:** Movement of a SUN-1-mRuby patch and its trajectory in a *mad-1* mutant nucleus. Movie is displayed at 10 frames per seconds.

**Video 3:** Movement of a SUN-1-mRuby patch and its trajectory in a *bub-3* mutant nucleus. Movie is displayed at 10 frames per seconds.

## Notes

### Competing Interest Statement

The authors have declared no competing interest.

